# The MacqD Deep Learning-based Model for Automatic Detection of Socially Housed Laboratory Macaques

**DOI:** 10.1101/2024.12.23.629644

**Authors:** Genevieve Jiawei Moat, Maxime Gaudet-Trafit, Julian Paul, Jaume Bacardit, Suliann Ben Hamed, Colline Poirier

## Abstract

Despite advancements in video-based behaviour analysis and detection models for various species, existing methods are suboptimal to detect macaques in complex laboratory environments. To address this gap, we present MacqD, a modified Mask R-CNN model incorporating a SWIN transformer backbone for enhanced attention-based feature extraction. MacqD robustly detects macaques in their home-cage under challenging scenarios, including occlusions, glass reflections, and overexposure to light. To evaluate MacqD and compare its performance against pre-existing macaque detection models, we collected and analysed video frames from 20 caged rhesus macaques at Newcastle University, UK. Our results demonstrate MacqD’s superiority, achieving a median F1-score of 99% for frames with a single macaque in the focal cage (surpassing the next-best model by 21%) and 90% for frames with two macaques. Generalisation tests on frames from a different set of macaques from the same animal facility yielded median F1-scores of 95% for frames with a single macaque (surpassing the next-best model by 15%) and 81% for frames with two macaques (surpassing the alternative approach by 39%). Finally, MacqD was applied to videos of paired macaques from another facility and resulted in F1-score of 90%, reflecting its strong generalisation capacity. This study highlights MacqD’s effectiveness in accurately detecting macaques across diverse settings.

## Introduction

Monitoring animal behaviour is crucial for comprehending their welfare status^1^ and brain function^2^. Traditional methods for monitoring animal behaviour, such as on-site observation or manual video analysis, are labour-intensive, prone to observer bias, and restricted by scalability and consistency^3^. Recent advances in machine learning offer new solutions for the automation of animal behaviour analysis, improving efficiency and reducing bias^4,5^. These developments have led to successful applications in small laboratory animals like mice and flies^3,6–9^, and are now emerging as promising tools for non-human primates (NHPs)^10–16^. Mimicking human observer behaviour, automatic behaviour analysis tools first detect and localise animals in an image, then classify the behaviours displayed. Accurate detection is therefore crucial, as errors at this stage lead to tracking failures and behaviour misclassification^17–19^.

Three main approaches have been used to detect animals in images or video recordings: (1) Background elimination, a non-deep learning approach that does not require training^20,21^; (2) Markerless keypoint estimation, which uses deep learning to detect and track animals based on anatomical landmarks (e.g., joints, eyes, ears)^22–29^; and (3) Deep-learning-based object detection, which utilises bounding boxes or pixel-level masks^19^. While the first two approaches are known to struggle when parts of the animal are occluded by an object or another individual^29,30^, a situation typical of complex environments and/or social settings, the last one is more resilient to this problem^18,19,31,32^.

NHPs serve as crucial models for understanding human cognition, neurobiology, and neuropathology due to their biological and cognitive similarities to humans^33–35^. Among them, macaques (Macaca sp.) are the most commonly used NHP model in biomedical research^36,37^. Monitoring their behaviour is essential in neuroscience, where they contribute to psychiatry^38,39^ and neurodevelopmental^40,41^ studies, and in animal welfare research aimed at reducing stress-related behaviours and improving housing conditions^42,43^. Compared to small laboratory animals like mice and flies, detecting macaques presents unique challenges due to their flexible joints, diverse postures, and lack of distinctive fur patterns in overlapping scenes.

A small number of studies have explored the automatic detection of macaques from video footage. Some studies focus on face detection for individual identification^44,45^, while others have attempted to detect the whole body of the animals for behaviour classification^17,46–50^. In terms of methods, most studies have used the general approaches described earlier (background extraction and deep learning methods). These tools typically achieve good macaque detection performance in simple settings but often fail when macaques are partially occluded. One exception is SIPEC^51^, a tool that integrates macaque detection, individual identification and behaviour classification. Using a deep-learning-based approach with pixel-level masks, SIPEC achieves state-of-the-art performance in macaque detection, including in scenarios of partial occlusion. However, its inference time is extremely long, and its generalisation performance (i.e. on individuals not seen during training) remains undocumented.

To address these challenges, we introduced MacqD, a Mask R-CNN-based model specifically designed to detect rhesus macaques in complex laboratory cages from video footage recorded using a single camera. We evaluated MacqD through a series of experiments comparing its robustness with pre-existing models, including SIPEC. After training the models (when appropriate) on images of specific individuals alone (Experiment 1) or in pairs (Experiment 2) in their cages, we evaluated their performance on new data from the same animals and on data from different animals. We then tested whether incorporating a tracking algorithm improves detection accuracy (Experiment 3). Finally, we further assessed MacqD’s ability to generalise by testing it on footage featuring paired macaques from a different animal facility (Experiment 4). Together, these experiments highlight the unique strengths of MacqD, namely, (1) its ability to deal effectively with occlusions, including overlapping macaque bodies, in challenging conditions (e.g. light over-exposure; glass reflection); and (2) its strong ability to generalise to videos from individuals and research facilities not used for training.

## Materials and Methods

### Data Collection

A collection of video recordings, subsequently referred as *Macaque* data, was acquired at the macaque research facility of Newcastle University, UK, between 2014 and 2020. The facility complies with the NC3Rs Guidelines for ‘Primate Accommodation, care and use’^52^, and comprises cages 2.1 m wide, 3 m deep, and 2.4 m high, exceeding the minimal requirement of the UK legislation, and where animals are housed in pairs. Besides the presence of a social partner, the cages are enriched by a multitude of structural elements and objects (e.g. shelves, swings, ropes) to promote the well-being of the animals. Data recording was approved by Newcastle University Animal Welfare and Ethical Review Body (project number: ID 928).

Video recordings were collected from 20 macaques which were selected as the primary subjects of observation (focal), using a remotely-monitored, wall-mounted digital cameras (Cube HD Y-cam, 1080p and Axis M1065-L, 1080p), fixed outside and positioned directly opposite each focal cage. While the cameras remained stationary for most of the study, they could be manually repositioned or zoomed in/out when necessary to improve visibility. Data were stored in .mp4 or .mov formats, with a spatial resolution of 1280 × 720 pixels, and sampled at 15 frames per second. The number of macaques visible on videos was variable. While by default the two cagemates were present, one animal was sometimes temporarily absent (e.g. when it was in the experimental laboratory). Due to the positioning of some cages back-to back, some videos also included non-focal animals in neighbouring cages. Examples of video frames from *Macaque* data are shown in Figure 1.

**Figure 1.**
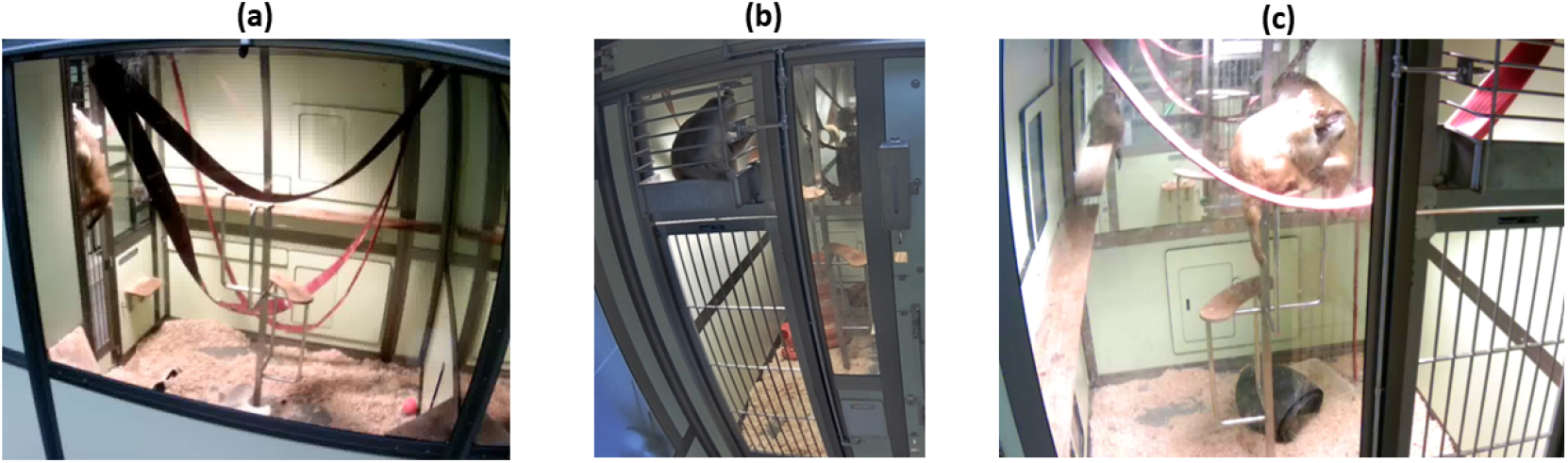
Example video frames used in this study. (**a**) Single macaque, partially hidden, with light overexposure;(**b**) Single macaque with cage railing occlusion; and (**c**) Pair of macaques in the focal cage, with partial overlap of the two individuals, partial occlusion from cage enrichment and one macaque from a neighbouring cage appearing in the background.

To further evaluate the model’s generalisation capabilities, a video from the Institut des Sciences Cognitives Marc Jeannerod, referred to as the ISC dataset, was also used. The ISC dataset has a frame rate of 24 frames per second and a spatial resolution of 2880 × 2160 pixels.

### Data Description

*Macaque* data were divided in several training and testing datasets (see Table 1) to be used in Experiments 1 and 2. Experiments 1 and 2 only differ by the number of macaques in the focal cage, respectively one and two. For the training datasets, video recordings from 10 individuals (either alone or in pairs) were selected. The same individuals were used for both experiments. Individual video frames were pseudo-randomly selected from recordings spanning various dates and times of day, ensuring that the datasets encompassed a wide range of cage settings, macaque postures, and positions within the cage. This approach aimed to ensure the training dataset were representative of the macaques housed at the Newcastle facility.

**Table 1.** Description of our *Macaque* data sub-setting into different training and testing datasets. For paired animals (Experiment 2), the same videos were used to assess the detection of each pair member (see **Results** section for details).

| Single or paired macaques | Training or test dataset | Same or different macaques as training dataset | Number of frames |
| --- | --- | --- | --- |
| Single | Training | - | 2,160 |
|  | Test | 'Same' | 45,000 |
|  |  | 'Different' | 45,000 |
| Paired | Training | - | 2,400 |
|  | Test | 'Same' | 22,500 |
|  |  | 'Different' | 22,500 |

For the testing datasets, five-minute videos from each single individual (Experiment 1) and each pair (Experiment 2) were used. In both experiments, two different testing datasets were employed: (1) the ‘Same’ dataset, comprising new video recordings of the same 10 individuals used in training, and (2) the ‘Different’ dataset, consisting of recordings from 10 new individuals. The ‘Different’ dataset was included to assess the generalisability of the models to macaques not encountered during training. While the training datasets were composed of isolated frames, we used consecutive frames for testing, in order to mirror real-world scenarios where behaviour recognition applications require dynamic information across frames. Because consecutive frames are often highly similar in terms of macaque cage position and body posture, the total number of frames was increased by a factor of 20 to enhance variability and representation. As a result, each video in the testing datasets consisted of 45,000 frames for videos featuring single macaques and 22,500 frames for paired macaques (see Table 1). Further variability was ensured by using videos from a relatively high number of individuals (n = 10).

Both the training and testing datasets included instances of occlusion, reflections, rapid motion, and overexposure. They also incorporated footage of animals displaying a comprehensive list of natural macaque behaviours (e.g. slow and fast locomotion, body shaking, foraging, interacting with objects, allogrooming and self-scratching). These diverse challenges were incorporated to enhance the comprehensiveness of the study and ensure that the analysis was conducted under realistic and varied conditions.

Additionally, a 17-second video from the Institut des Sciences Cognitives Marc Jeannerod (ISC dataset) containing 420 frames was used to further test the model. This video presented additional challenges, including occlusion, the presence of objects such as toys used for stimuli, and human reflections on the glass.

### Data Annotation

Training and testing datasets were annotated using the VIA image annotator^53^. In the training dataset, each video frame was annotated with pixel-level masks, a technique known as segmentation, where each pixel is assigned to an individual macaque. For the testing datasets, including the ISC dataset, annotations were made with bounding boxes, with rectangles drawn around each macaque to include all body parts while minimising the box area. This approach was selected to facilitate the comparison of different algorithms, some of which only output bounding boxes (see subsection **Performance Metrics**). Annotations were performed by ten different research assistants, with each annotation verified by at least one other assistant and subsequently validated by the first authors. In the training datasets, macaques visible in neighbouring cages were also annotated to maximise learning, whereas for the testing datasets, the detections and ground truths of macaques in the background were excluded to focus on evaluating how well models detected animals in the focal cage. This approach does not count correct detections of neighbouring macaques toward model performance and does not penalise the model for failing to detect them.

### Macaque Detection Algorithms

In this study, macaque detection was assessed using three different algorithms. The first two implemented a deep learning approach using Mask R-CNN as the framework, training a neural network through supervised learning (where the network learns from labelled examples). The third algorithm was based on background elimination, an approach that does not require any training.

#### Mask R-CNN

Mask R-CNN^54^ is a state-of-the-art, two-stage framework widely used for segmenting individual objects in images (instance segmentation). It not only detects objects but also outlines their precise location with pixel-level masks. In the first stage, the image passes through a feature extractor (a series of convolutional layers) that generate feature maps representing key characteristics of the image. These maps are enhanced by a Feature Pyramid Network (FPN), which improves detection at different scales by combining detailed high-resolution features with more abstract low-resolution ones. This helps the model detect objects of various sizes in complex scenes. The refined maps are then processed by the Region Proposal Network (RPN), which suggests areas likely to contain objects (Regions of Interest, or ROIs). In the second stage, features from each ROI are extracted using RoIAlign, a method that ensures precise alignment with the original image. This alignment is critical for refining bounding boxes, classifying objects, and generating accurate segmentation masks, especially for small or detailed objects (Fig. 2).

**Figure 2.**
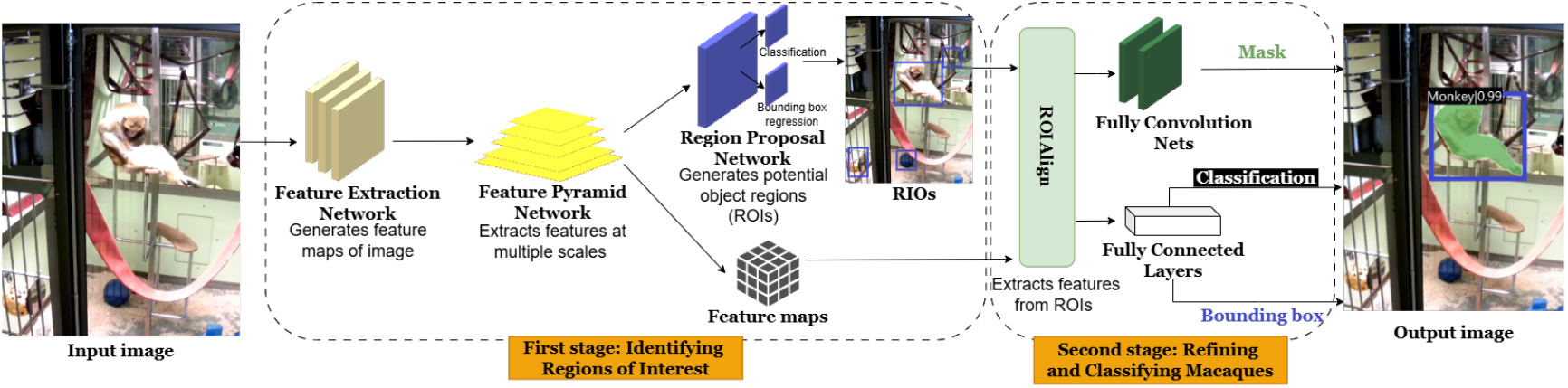
Overview of the Mask R-CNN framework for macaque detection.

In this study, a modified Mask R-CNN model, referred to as MacqD, was created for detecting macaques in video frames, incorporating SWIN^55^, a state-of-the-art transformer-based feature extractor. Throughout this paper, the term ‘MacqD’ refers specifically to this modified model. The images processed in MacqD were resized to a maximum of 1333 × 800 pixels (smaller images retained their original size), maintaining the aspect ratio. Images were padded to meet network requirements and normalised using standard ImageNet values. During training, random horizontal flipping was applied to improve the model’s generalisability by increasing dataset variability while preserving biological validity, while no changes were applied during testing to ensure consistent evaluation. As a benchmark, MacqD was compared with SegNet, another Mask R-CNN variant implemented in SIPEC^51^, which uses ResNet101^56^, a widely used convolutional neural network, as its feature extractor. SegNet was trained on frames resized to 1280 × 1280 pixels, maintaining the aspect ratio. During training, random rectangular regions were hidden (converted to black pixels) to help detect partially occluded objects, while no data augmentation was applied during testing.

#### Background Elimination (BE)

Unlike deep-learning techniques which require extensive training and computational resources, the background elimination method is more efficient, as it does not require any training. In this research, an optimised Background Elimination (BE) pipeline was designed specifically for extracting macaques from video footage.

Our pipeline (Fig. 3) processes a video by extracting every 10th frame to build a background image, which is first resized to 1280 × 720 pixels to ensure consistency. The extracted frames are grouped into sets of 120, slightly blurred to smooth out minor variations, and the 70th percentile of each pixel’s RGB (red, green, blue) values is calculated to create sub-background images. These sub-backgrounds are then combined by taking the median RGB values to generate the final background image. Each video frame is compared to this background using the MOG2^57^ algorithm from the OpenCV library^58^, which detects macaques by separating them from the static background. Even after the background is removed, small amounts of noise or gaps within the macaque’s outline may remain. To correct these imperfections, morphological operations are applied to group nearby pixels into clusters and fill gaps. A convex hull is then calculated for each cluster, forming a polygon that encloses the object by connecting its outermost points. The overlapping convex hulls are merged to smooth the outline, and small irrelevant clusters are filtered out, isolating the primary macaque. Finally, a bounding box is placed around the refined cluster to localise the detected macaque within the frame.

**Figure 3.**
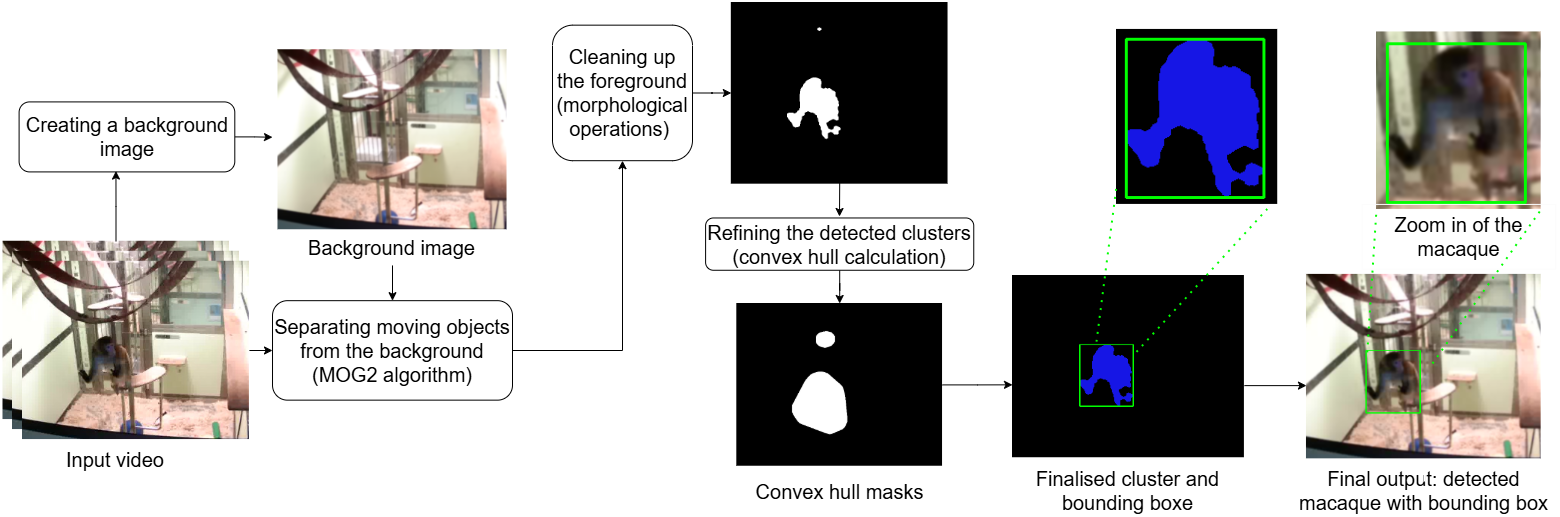
Overview of the background elimination pipeline.

### Experimental Design - Experiments 1 & 2

Figure 4 illustrates the models used in the two first experiments: Experiment 1 tested models on single macaque recognition, and Experiment 2 on paired macaques. In Experiment 1, the MacqD model was trained on a dataset where a single animal was present in the focal cage (*Macaque Single* dataset) and compared with background elimination (BE) and three SegNet variants: SegNet - Primate, from the original paper^51^, trained on a primate dataset with macaque images from the authors’ research facility; SegNet - Macaque Single, trained exclusively on our *Macaque Single* dataset; and SegNet - Primate + Macaque Single, which used SegNet - Primate as a starting point and was further trained with our *Macaque Single* dataset.

**Figure 4.**
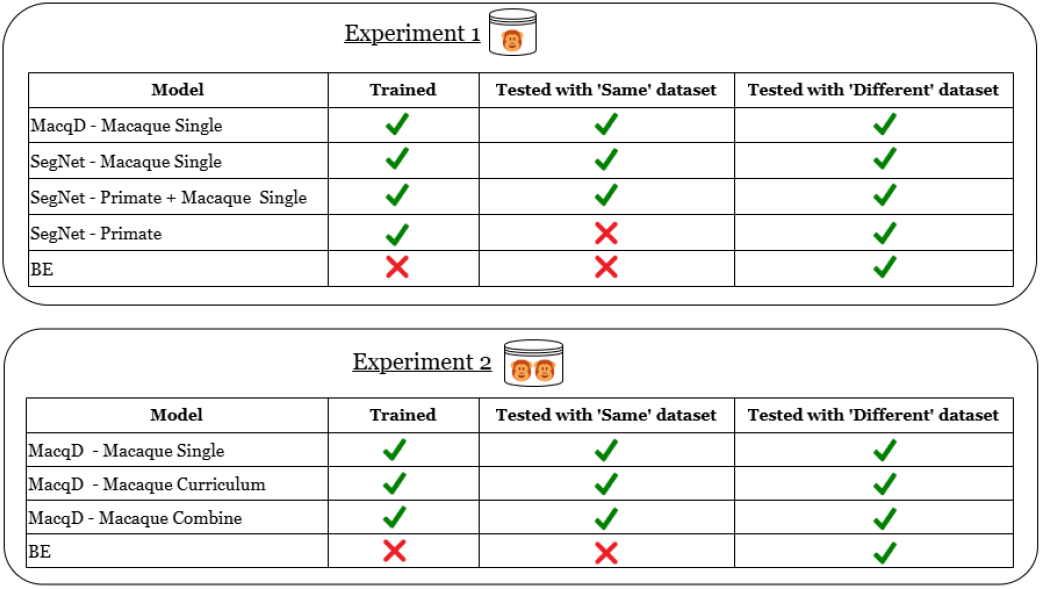
Overview of experiments 1 and 2 illustrating how the models compared in this study differed in terms of training and testing datasets. ‘Same’ and ‘Different’ correspond to the datasets described in Table 1, with the labels referring to the fact that the model was tested with videos of individual macaques ‘seen’ during the training phase (‘same’) or not (‘different’) (see subsection **Data Description** for more details).

In Experiment 2, the MacqD model trained on the *Macaque Single* dataset was compared to two other MacqD derivatives: MacqD - Macaque Curriculum, which used MacqD - Macaque Single as a starting point and was further trained with the dataset featuring paired macaques in the focal cage (*Macaque Paired*); and MacqD - Macaque Combine, which was trained on a merged training dataset combining *Macaque Single* and *Macaque Paired* datasets. MacqD - Macaque Curriculum was used to assess curriculum learning^59^, a strategy where models are trained by gradually increasing task complexity, mimicking human learning by starting with simpler concepts and progressing to more difficult ones. In contrast, MacqD - Macaque Combine was trained on a combined dataset, exposing the model to diverse scenarios all at once while saving time. MacqD models were also compared to BE but not SegNet models, due to poor results obtained with these models in Experiment 1 (see subsection **Experiment 1: Detection of Single Macaques**). All MacqD models and the SegNet - Macaque Single model were trained for 100 epochs, with the final model selected based on the epoch with the minimum validation loss. All final models were tested on the *Macaque* ‘different’ dataset and, where applicable, the *Macaque* ‘same’ datasets (Fig. 4).

### Tracking Algorithm (Experiment 3)

In computer vision, tracking algorithms monitor object movement across consecutive video frames by estimating the target object’s positions in subsequent frames, given its initialised position^60,61^. In Experiment 3, results from the detection models were compared before and after implementing a tracking algorithm to test performance improvement. The tracking algorithm aims to maintain detection continuity across frames by estimating the location and motion of macaques, thereby reducing instances where the macaque is not detected in subsequent frames.

The centroid (geometric centre) of each detected macaque, whether from a mask produced by MacqD and SegNet or a cluster from BE, was used as input for the Kalman filter^62^, a mathematical algorithm that predicts an object’s position based on past movements. If a macaque was not detected in a given frame, the Kalman filter estimated its position using the centroid from previous frames while retaining the same bounding box size from the last known detection. This prediction process continued until a match was found between the predicted and detected positions or until 20 frames had passed without a match. To associate predicted positions with current detections, the Hungarian algorithm^63^ was used. This algorithm matches the predicted position of an object to the closest detected object in the current frame, identifying which detection belongs to which macaque (see Fig. 5).

**Figure 5.**
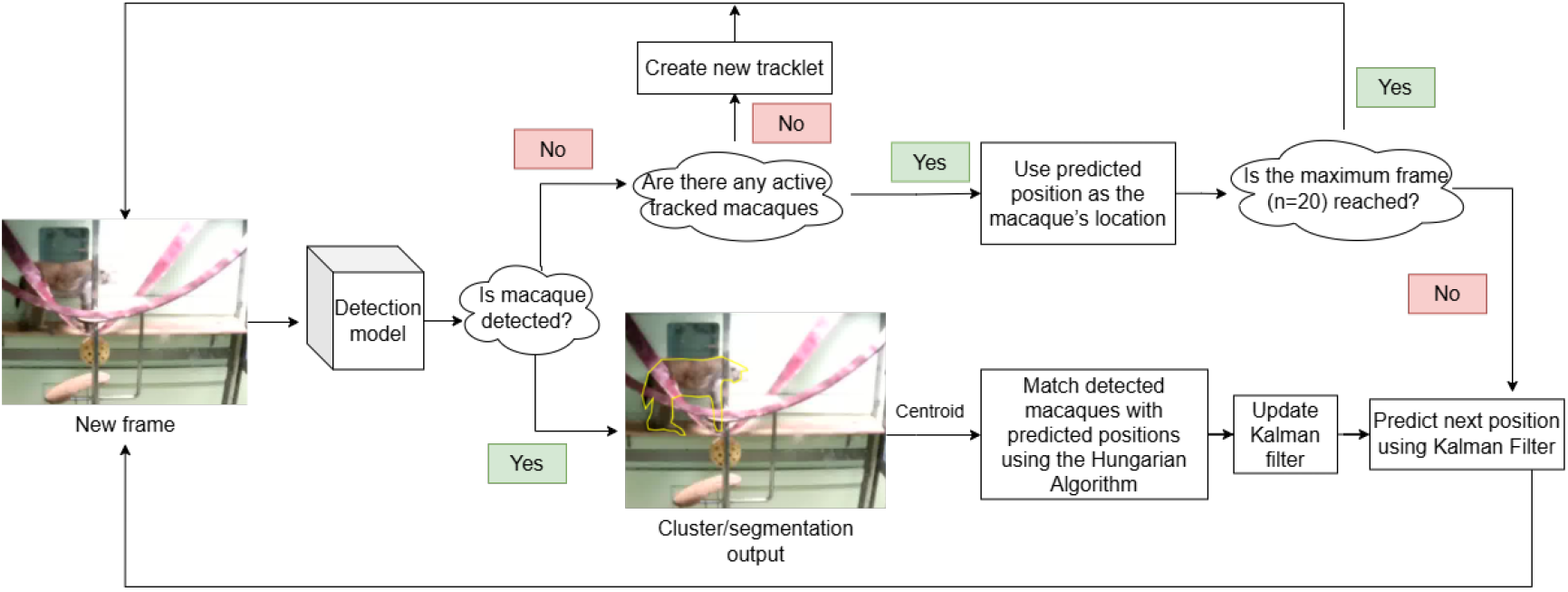
Tracking algorithm pipeline.

### Performance Metrics

Evaluation of the different models was based on bounding boxes in order to standardise comparisons across all models (MacqD and SegNet provide bounding boxes and pixel-based masks but BE only outputs bounding boxes). The Intersection over Union (IoU) metric was used to measure the overlap between predicted and ground truth boxes, with an IoU of 0.50 or higher considered a true positive (TP) and an IoU below this threshold classified as a false positive (FP).

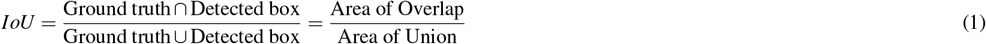

Performance was assessed using precision, recall, and the F1 score applied at the individual level, with precision measuring how often the model was correct when it identified a portion of the frame as containing an individual macaque, recall measuring the model’s ability to not miss a macaque when this one was present in the frame. The F1 score is the harmonic mean of precision and recall. It ensures that F1 is high only when both precision and recall are high (for instance reaching 1 only if both are 1), and low when both are low (dropping to 0 if either is 0). Macaques in the non-focal cage were ignored.

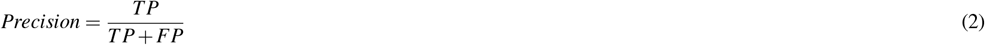

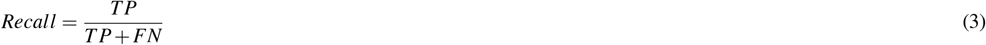

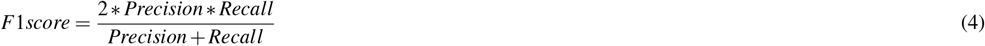

MacqD and SegNet models provide a confidence score to each bounding box, indicating the likelihood of correct identification. In Experiment 1, the optimal threshold for filtering out false positives was determined by maximising precision and recall performance metrics based on the validation dataset. Confidence scores were assessed from 0.50 to 0.95 in 0.05 increments, and the median precision vs. recall was plotted to identify the optimal threshold (see Supplementary material, Fig. S1).

### Statistical Test

Precision, recall, and F1 score were used for statistical tests to assess the differences in performance. Initial attempts to fit linear mixed-effects models indicated non-normally distributed residuals, violating parametric assumptions. Consequently, non-parametric approaches previously applied for evaluating machine learning methods were utilised^64^. The Friedman test compared the performance of multiple models across datasets, while the Wilcoxon (signed-rank) test assessed the performance of pairs of models. The Benjamini-Hochberg procedure^65^ was employed to control the false discovery rate for multiple comparisons. Additionally, Wilcoxon tests were conducted to compare results before and after implementing the tracking algorithm.

### Computing Environment

MacqD models were implemented from the open-source framework MMDetection version 2.25^66^, and SegNet models from the open-source pipeline^67^. We used Python (3.7.13), CUDA toolkit (10.1) and GPU-accelerated library (CUDNN 7.6.3) with NVIDIA GeForce GTX 1080 Ti GPU. Python (3.6.13), CUDA (10.2.89) and CUDNN (7.6.5) were used to develop BE.

## Results

### Experiment 1: Detection of Single Macaques

Comparing models tested with new video recordings of the 10 single individuals used for training (‘Same’ dataset), a Friedman test revealed significant differences in precision, recall, and F1 score among the models (Fig.6 and Table 2). The performance of MacqD - Macaque Single was particularly high, with a median precision of 0.99, recall of 0.99, and F1 score of 0.99 (Table 2, see Supplementary Material Fig. S2a for additional evaluation on different durations). Wilcoxon pair-wise tests revealed that MacqD - Macaque Single was significantly better than other models for the three metrics (Table 2).

**Table 2.**
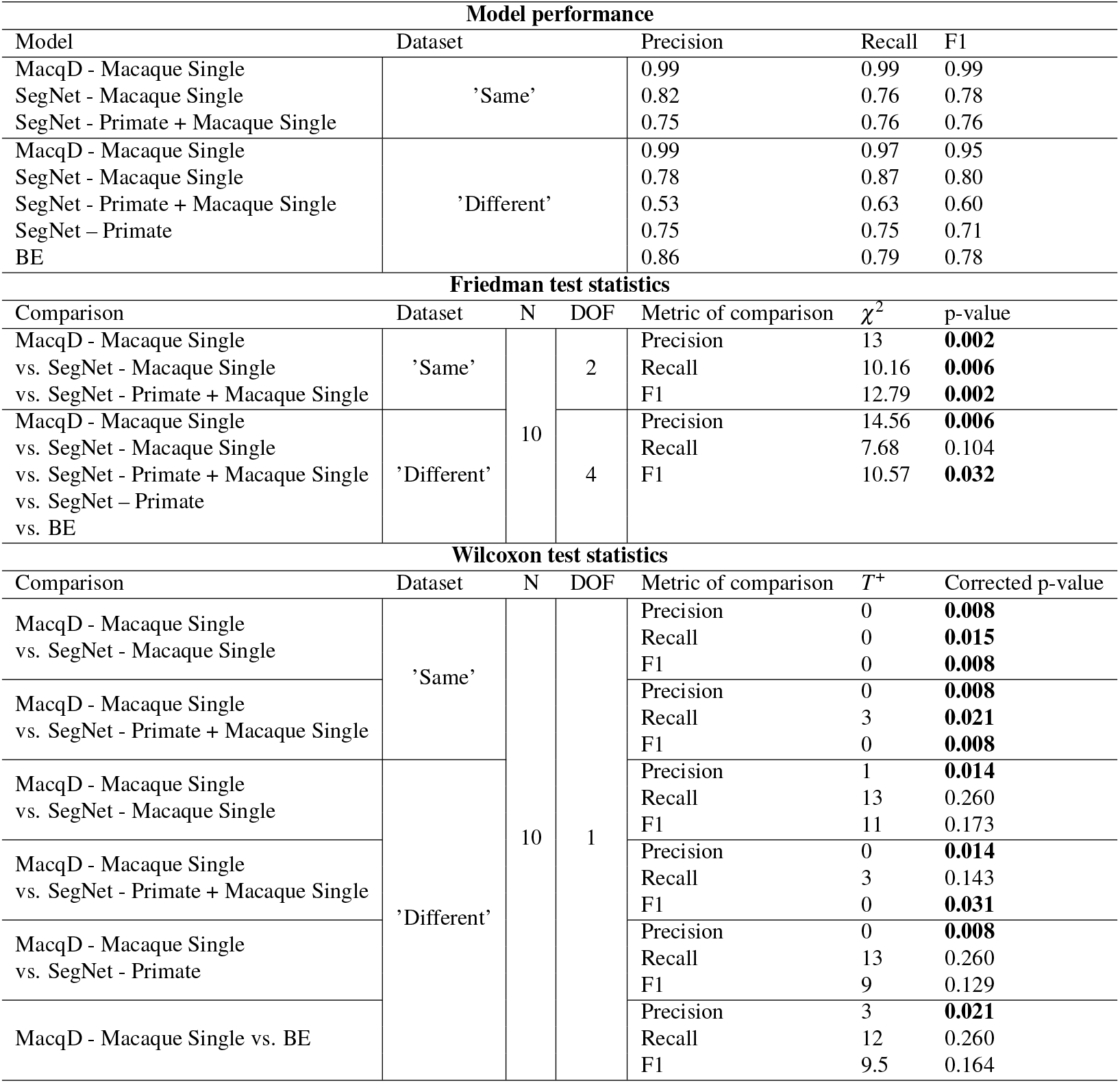
Model performance and statistical comparisons of video featuring single macaques (Experiment 1). Bolded p-values indicate significant at p<0.05.

In evaluating videos with animals not included in the training dataset (‘Different’ dataset), the Friedman test indicated significant differences among the models in precision and F1 score, but not in recall (Fig. 6 and Table 2). MacqD - Macaque Single also demonstrated notably high performance, achieving a median precision of 0.99, recall of 0.97 and F1 score of 0.95 (Table 2, see Supplementary Material Fig. S2a for additional evaluation on different durations). Wilcoxon tests revealed that MacqD - Macaque Single was significantly better than any other tested model in precision and significantly better than SegNet - Primate + Macaque Single in F1 score (Table 2). The two first columns of Fig. 8 visually illustrate MacqD’s generalisation and detection capabilities, in particularly challenging scenarios, further supporting these results.

**Figure 6.**
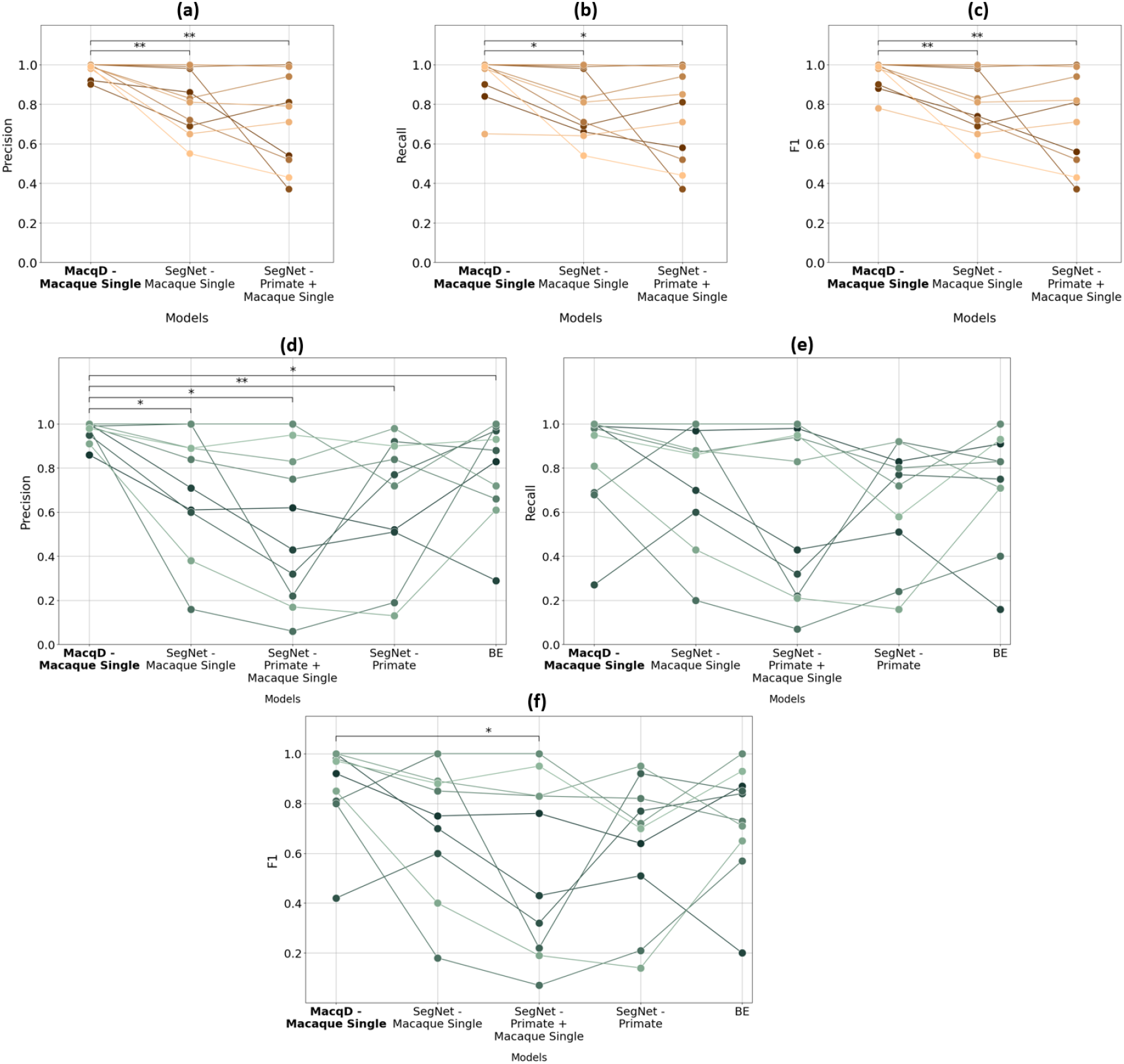
Model performance in Experiment 1, where the test datasets contain only a single macaque in the focal cage. (**a**), (**b**), and (**c**) represent precision, recall, and F1 scores, respectively, evaluated on datasets featuring the same macaques (but different videos) as the training dataset (‘Same’ dataset). (**d**), (**e**), and (**f**) represent precision, recall, and F1 scores, respectively, evaluated on datasets featuring macaques not presenting in the training dataset (‘Different’ dataset). Markers represent results for individual macaques, with lines connecting the markers across models to illustrate performance variations. Wilcoxon test: *p<0.05, **p<0.01.

In addition to performance metrics, training and testing times are crucial parameters for evaluating and comparing models. SegNet models required the longest training time (10 hours for 100 epochs) and inference time (3 days 10 hours hours for a 5-minute video), followed by MacqD models (train: 9 hours for 100 epochs; inference: 17 minutes for a 5-minute video). BE was the fastest overall, requiring only 4 minutes for inference for a 5-minute video. Considering both performance and time factors, only MacqD and BE were used in the subsequent experiment to analyse the paired macaque datasets.

### Experiment 2: Detection of Paired Macaques

Comparing models tested with new video recordings of the 10 paired individuals used for training (‘Same’ dataset), a Friedman test revealed significant differences in precision, recall, and F1 score among the models (see Fig. 7 and Table 3). MacqD - Macaque Curriculum and MacqD - Macaque Combine achieved similar results, with median F1 scores of 0.9 and 0.87, respectively. Among these models, MacqD - Macaque Curriculum had the highest recall of 0.87, while MacqD - Macaque Combine achieved the highest precision of 0.97 (see Table 3, see Supplementary Material Fig. S2b, S2c and S2d for additional evaluation on different durations). Wilcoxon pairwise tests indicated that MacqD - Macaque Curriculum and MacqD - Macaque Combine were both significantly better than MacqD - Macaque Single in precision and F1 score (Table 3).

**Table 3.**
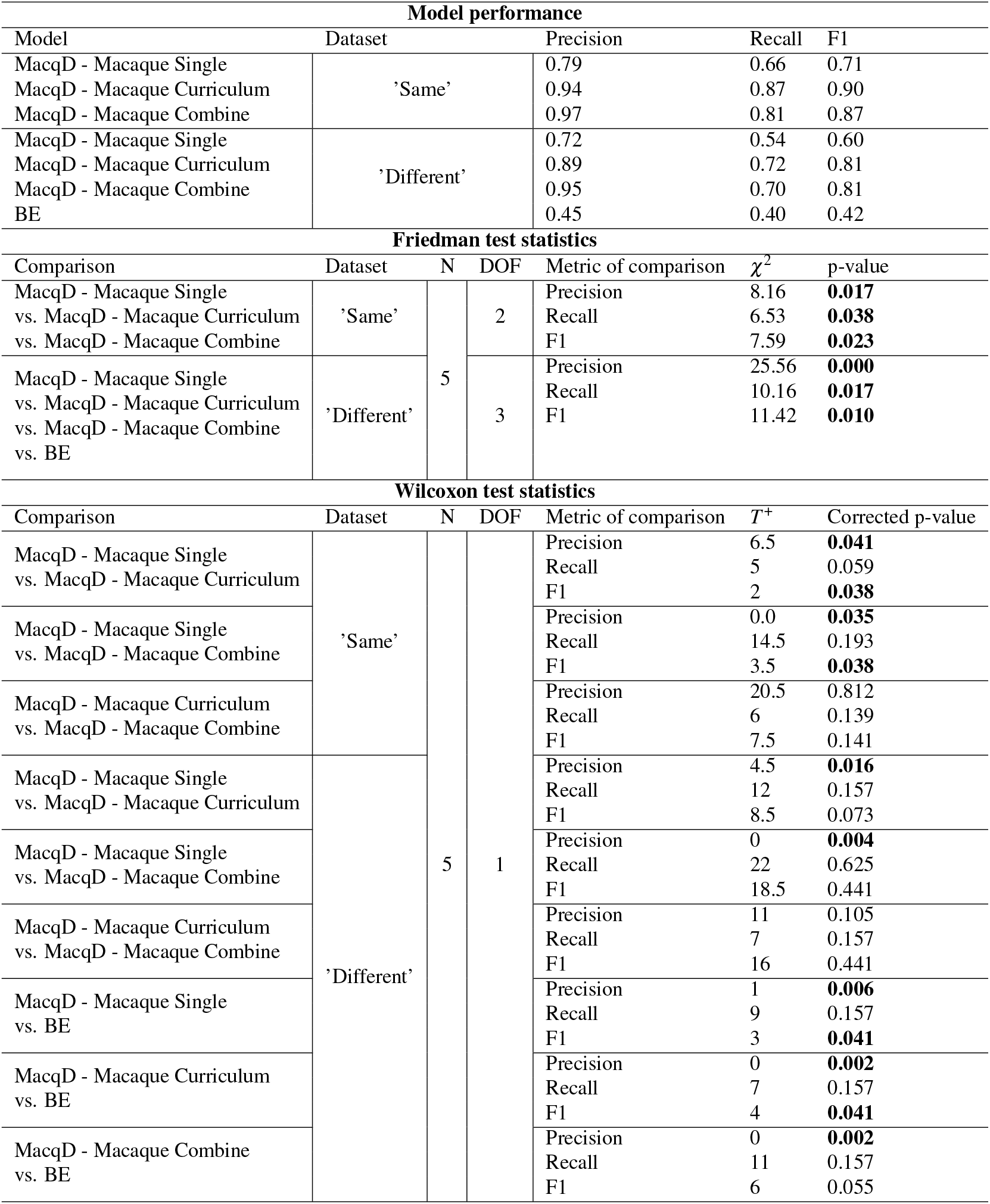
Model performance and statistical comparisons of video featuring paired macaques (Experiment 2). Bolded p-values indicate significant at p<0.05.

**Figure 7.**
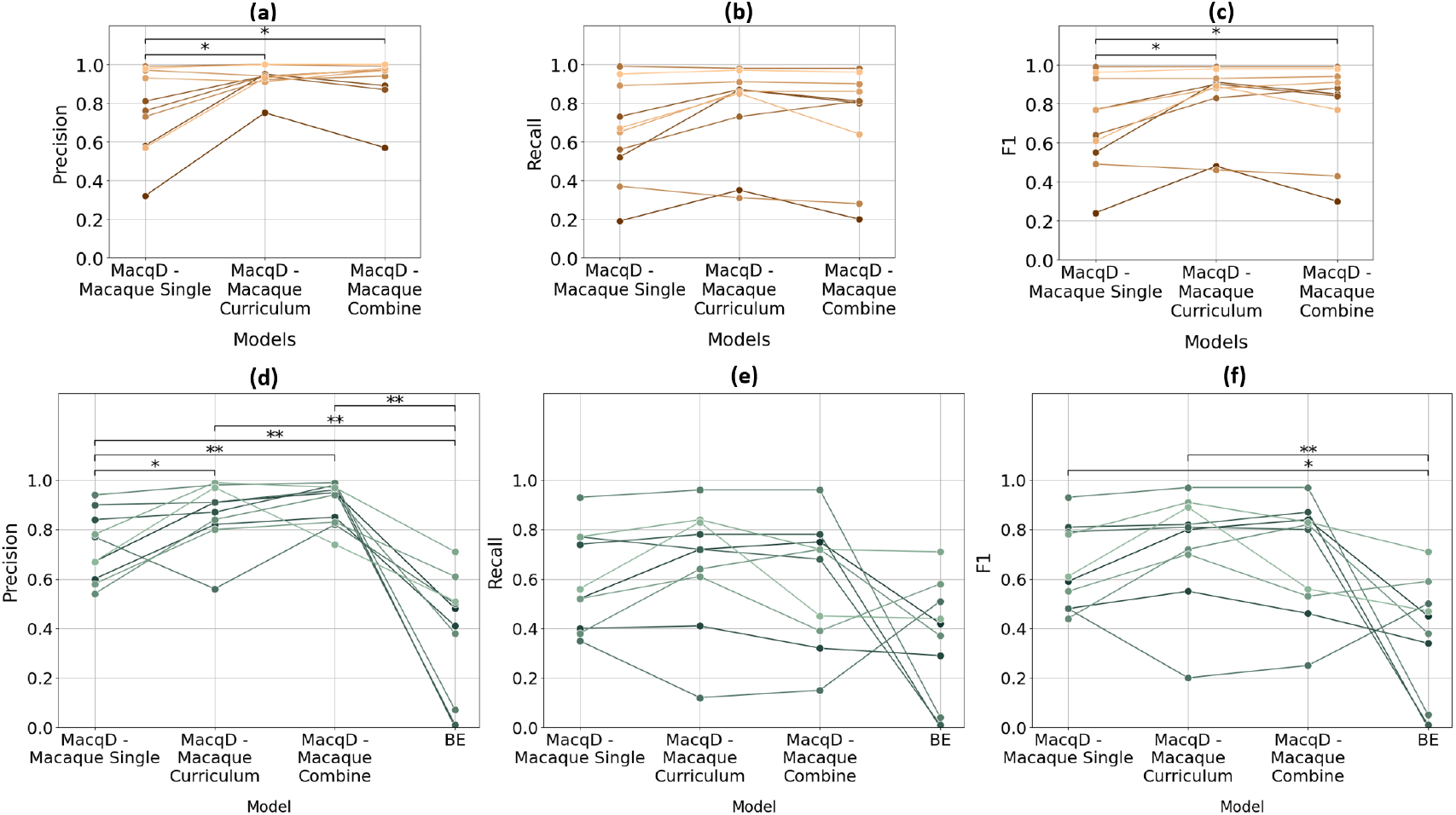
Model performance in Experiment 2, where the test datasets contain paired macaques in the focal cage. (**a**), (**b**), and (**c**) represent precision, recall, and F1 scores, respectively, evaluated on datasets featuring the same macaques (but different videos) as the training dataset (‘Same’ dataset). (**d**), (**e**), and (**f**) represent precision, recall, and F1 scores, respectively, evaluated on datasets featuring macaques not included in the training dataset (‘Different’ dataset). Markers represent results for individual macaques, with lines connecting the markers across models to illustrate performance variations. Wilcoxon test: *p<0.05, **p<0.01.

In evaluating videos featuring different animals than those featured in the training dataset (‘Different’ dataset), the Friedman test indicated significant differences among the models in precision, recall, and F1 score (Fig. 7 and Table 3). MacqD - Macaque Curriculum and MacqD - Macaque Combine achieved very similar results, with median F1 scores of 0.81. MacqD - Macaque Curriculum had the highest recall of 0.72, while MacqD - Macaque Combine achieved the highest precision of 0.95 (Table 3, see Supplementary Material Fig. S2b, S2c and S2d for additional evaluation on different durations). Wilcoxon pairwise tests revealed that MacqD - Macaque Curriculum and MacqD - Macaque Combine were both significantly better than MacqD - Macaque Single in precision (Table 3). Additionally, all other models significantly outperformed BE in precision. For the F1 score, BE was significantly outperformed by MacqD - Single and MacqD - Macaque Curriculum. The last two columns of Fig. 8 further illustrate MacqD’s generalisation capabilities, particularly in challenging scenarios where macaques overlap in the video.

**Figure 8.**
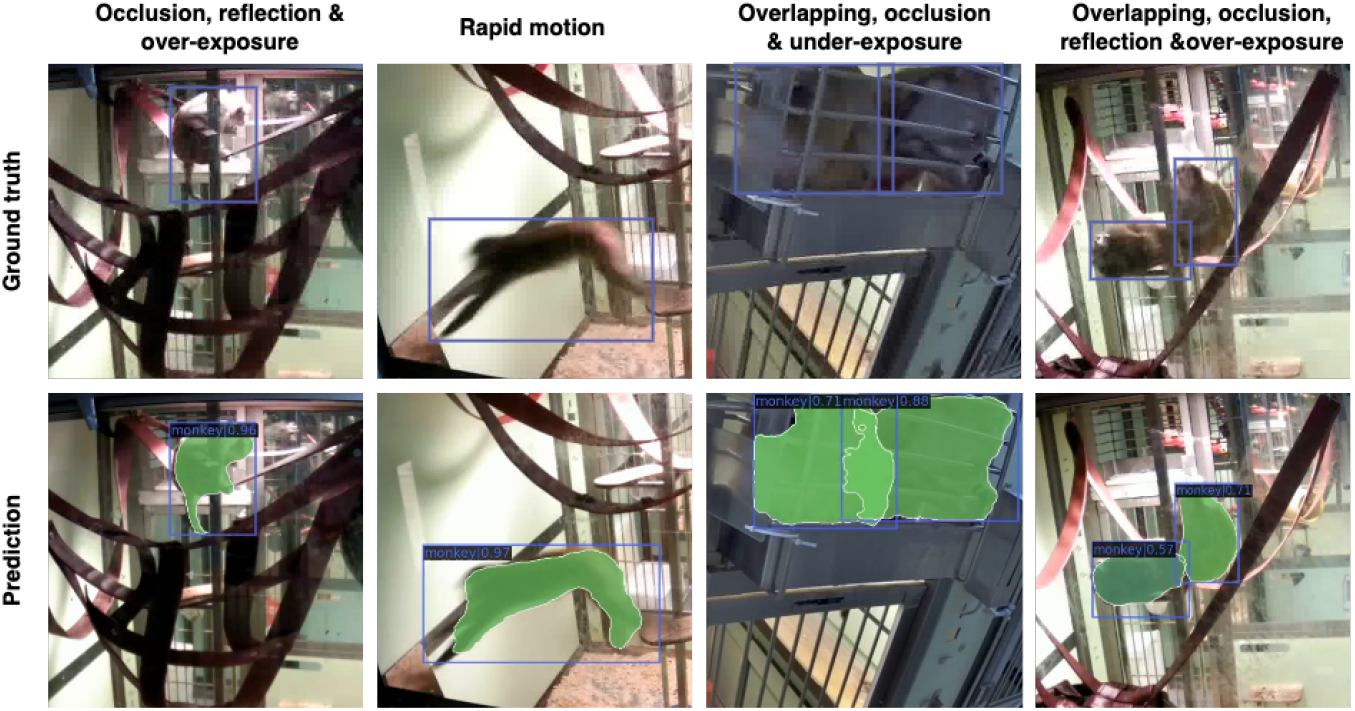
Comparison of ground truth versus predicted segmentations/bounding boxes for frames from unseen animals. The first two columns show MacqD - Macaque Single’s predictions on frames featuring single animals not present in the training dataset (Experiment 1, ‘Different’ dataset). The third and fourth columns show MacqD - Macaque Combine’s predictions on frames featuring pairs of macaques not present in the training dataset (Experiment 2, ‘Different’ dataset).

Although the Friedman test revealed significant differences between the models in recall, the Wilcoxon pairwise tests did not find any significant differences in recall after p-value correction for both testing the ‘Same’ and ‘Different’ datasets. There were no significant difference between MacqD - Macaque Curriculum and MacqD - Macaque Combine in any of the metrics.

### Experiment 3: Impact of Tracking Algorithm on Detection Performance

To evaluate potential improvements in model performance, we hypothesised that applying a tracking algorithm could enhance recall by reducing missed detections between frames. When comparing detection results before and after applying the tracking algorithm on the dataset featuring single macaques (Experiment 1) that were present in the training dataset (‘Same’ dataset), the Wilcoxon pairwise test revealed no significant differences in any metrics (see Supplementary Material, Fig. S3). When comparing detection results before and after applying the tracking algorithm on the dataset featuring macaques that were not present in the training dataset (‘Different’ dataset), the Wilcoxon pairwise test found a significant increase in recall for the BE model only, with recall improving from 0.79 to 0.82 (see Supplementary Material, Fig. S4 and Table S1).

In contrast, when comparing detection results before and after applying the tracking algorithm on the dataset featuring paired macaques (Experiment 2) that were present in the training dataset (‘Same’ dataset), the Wilcoxon pairwise test revealed significant decreases in precision for both the MacqD - Macaque Single and MacqD - Macaque Combine models (see Supplementary Material Fig. S5). These minor decreases (1-2%) suggest that while tracking may improve recall (see Supplementary Material Table S2), it can also reduce precision, likely due to increased false positives (see **Discussion** section for details). For performance tested on the dataset that included macaques not present in the training data (‘Different’ dataset), no significant differences in any metrics were observed before and after applying the tracking algorithm (see Supplementary Material, Fig. S6 and Table S2).

### Experiment 4: performance on ISC dataset

To further evaluate the generalisation of the best models, we tested MacqD - Macaque Curriculum and MacqD - Macaque Combine on a video from another research facility (ISC dataset) featuring paired macaques. As shown in Table 4, both models achieved the same F1 score of 0.87. After applying the tracking algorithm, MacqD - Macaque Curriculum’s F1 score improved to 0.90.

**Table 4.** Results of ISC dataset evaluation.

| Model | Before tracking implementation |  |  | After tracking implementation |  |  |
| --- | --- | --- | --- | --- | --- | --- |
|  | Precision | Recall | F1 | Precision | Recall | F1 |
| MacqD - Macaque Curriculum | 0.89 | 0.86 | 0.87 | 0.91 | 0.89 | 0.90 |
| MacqD - Macaque Combine | 0.92 | 0.82 | 0.87 | 0.90 | 0.82 | 0.86 |

## Discussion

This study aimed to develop a robust tool for detecting macaques under challenging laboratory conditions, which led to the creation of MacqD. We demonstrated MacqD’s ability to detect both single and paired macaques accurately, even in scenarios involving occlusions, glass reflections, and overexposure. MacqD was tested on an extensive dataset of 90,000 images, far surpassing SegNet’s 191 frames^51^. Compared to SegNet^51^, MacqD was tested on unseen individuals as well as footage from a different facility, highlighting its robust and generalisable performance. To address the challenges of occlusion, caused either by objects or overlapping individuals, as commonly encountered in animal detection^19,47,68^, we conducted a stepwise evaluation of MacqD’s performance under different occlusion conditions. This rigorous approach enables comparisons with other pre-existing models, highlighting MacqD’s strengths in handling these scenarios.

We first evaluated MacqD on videos containing single macaques, where occlusions were caused by cage structures or enrichment within the cage, while glass reflections and overexposure from lighting often obscured parts of the macaques. As a result, MacqD performed robustly under these conditions, accurately detecting macaques despite partial occlusions. In contrast, the BE model, while computationally efficient and requiring no training, struggled to detect static macaques, frequently misclassifying them as background. MacqD outperforms both SegNet models: one trained on its original dataset (SegNet - Primate) and the other with additional training using our data (SegNet - Primate + Macaque). This is likely due to their reliance on default parameters, as fine-tuning was computationally prohibitive. Overall, MacqD’s SWIN-based feature extraction proved more capable, yielding superior results than other pre-existing models in complex settings.

Overlapping macaques present specific challenges, as their similar fur patterns make distinguishing individuals difficult, even for human observers. To address this, we tested different versions of MacqD on videos featuring paired macaques and compared their performance with the BE model. SegNet was excluded due to its poor performance on single-macaque datasets. The BE model performed poorly, frequently merging closely positioned macaques into a single detection. In contrast, both MacqD - Macaque Curriculum and MacqD - Macaque Combine effectively detected individual macaques, demonstrating their ability to handle partially-overlapping subjects. There were only small differences between these two MacqD models, with MacqD - Curriculum slightly better in recall and MacqD - Combine slightly better for precision. However, MacqD trained only on video frames featuring single macaques (MacqD - Macaque Single) struggled in these scenarios, indicating that training solely on simple, non-overlapping images is inadequate for addressing such complexities. This limitation underscores the importance of a diverse training data.

To determine whether a five-minute test video efficiently represents MacqD’s performance, additional evaluations were conducted using shorter video segments (2, 3, and 4 minutes). The results (see Supplementary Material, Fig. S2) indicate stable F1 scores across all durations, confirming that five-minute evaluations are representative and reinforcing the robustness of the assessment approach.

In addition to evaluation duration, MacqD’s generalisation capabilities were assessed by testing it on data from different animals at Newcastle University and on a dataset from another facility (ISC dataset). Both MacqD - Macaque Curriculum and MacqD - Macaque Combine achieved good performance, demonstrating strong generalisation to new environments.

A tracking algorithm was applied to assess whether it would enhance detection results. Only small differences were observed, with both slight improvement and small deterioration, depending on the specific model, dataset and metric. The tracking algorithm tended to reduce false negatives by detecting missed macaques between consecutive frames, improving recall. However, at least in some cases, it also propagated false positives, where an incorrect detection in one frame was carried forward, lowering precision (for a counter-example, see results with MacqD - Curriculum on the ISC dataset in Fig. 4).

Despite its strong results, MacqD shares one limitation with existing tools. Like other deep learning-based models (but unlike background elimination approaches), MacqD cannot be used for real-time applications due to high computational demands. A partial solution is to record videos during daytime and process them overnight. Beyond this, while MacqD demonstrated the ability to generalise its good performance to videos from another research facility, its performance in different types of environments, such as those outdoor enclosures or recordings from moving cameras, remains uncertain and should be tested. Additionally, our evaluation focused on paired macaques. While we expect MacqD performance to be similar with slightly bigger groups (3-6 individuals), large macaque groups of ten or more individuals, typical of breeding centres, are likely to pose additional challenges. Future studies will need to test MacqD performance in these types of scenarios. Lastly, while the tracking algorithm improved recall in some datasets, it also increased false positives, reducing precision. Future implementations, such as advanced tracking techniques or hybrid approaches, could help strike a better balance between precision and recall, enhancing detection robustness in challenging scenarios.

With this publication, we release two versions of MacqD, MacqD - Curriculum and MacqD - Combine, along with a tracking algorithm, available here (https://github.com/C-Poirier-Lab/MacqD). For future users, we recommend MacqD - Curriculum to maximise overall performance or achieve the best recall, while MacqD - Combine is preferable for applications where precision is the primary metric. Both versions demonstrated strong generalisation on datasets from Newcastle University and ISC. However, while MacqD performs well on data from certain facilities (e.g., the ISC dataset), we cannot exclude the possibility that it will deliver suboptimal results in others. In such cases, we recommend further training MacqD with a small amount of locally collected data to improve performance. The tracking algorithm can enhance recall by reducing false negatives between consecutive frames, but it may also propagate false positives, lowering precision (see results from both models on ISC dataset; Fig. 4). We advise users to test the tracking algorithm on their own datasets, as its effects can vary.

Looking forward, MacqD can be used to automatically quantify the time an animal spends in specific parts of the cage. Such information can be useful for detecting when arboreal behaviour resumes after a surgical intervention or how often animals interact with enrichment elements^69^. Simple mathematical operations can also be applied directly to MacqD’s output to quantify how much, how fast, and when an animal is moving. This information could be used for monitoring purposes, automatically triggering alert messages that signal significant abrupt changes in habitual movement patterns. For socially housed animals, MacqD can be combined with a dedicated face detection model for individual identification^47^. However, we believe that the most exciting application of MacqD lies in combining it with a behaviour recognition algorithm, as a large amount of behavioural data could significantly enhance the scope of neuroscience and behavioural questions that can be addressed.

## Conclusion

In this study, we introduced MacqD, a Mask R-CNN-based model for detecting socially-housed macaques in indoor laboratory environments, using single-camera video footage. MacqD consistently demonstrated superior precision, recall, and F1 scores compared to pre-existing models, excelling in both single and paired macaque detection under challenging conditions. It also showcased strong generalisation to previously unseen individuals and new facilities, highlighting its adaptability and robustness. With its superior performance and low-cost setup, MacqD is a versatile tool for enhancing behavioural monitoring in neuroscience, animal welfare, and biomedical research

## Data Availability

The data used in this study, obtained from Newcastle University and the Institut des Sciences Cognitives Marc Jeannerod, contain sensitive records of non-human primates and are not publicly available. Data requests need to be discussed with the respective institutions. The source code for the algorithms used in this study is available in the GitHub repository https://github.com/C-Poirier-Lab/MacqD.

## Acknowledgements

We would like to thank Dr. J.Castlellano Bueno, G.Atkinson, M.Boddy, B.Banfield, L.Hannam, E.Hall, A.McKenna, R.Mishra, E.Stebbings, J.Tulip and S.Sanjeev for their help in acquiring and annotating the dataset for this study. Their hard work was a big part of making this research possible. This work was funded by Centre for Doctoral Training in Cloud Computing for Big Data [EP/L015358/1], the Barbour Foundation and a NC3Rs project grant [NC/K000802/1].

## Author Contributions Statement

Conceptualisation and Methodology: S.B.H., C.P., J.B.; Data Analysis: G.J.M., M.G-T.; Model Development: G.J.M., M.G-T., J.P., S.B.H., C.P.; Manuscript Preparation: G.J.M., M.G-T., S.B.H, J.B.; Supervision: S.B.H, J.B.,C.P.; Resource: S.B.H., J.P., C.P., J.B.; Visualisations and Figure: G.J.M.; Ethical Compliance and Approvals: C.P.

