## Supplemental Figure S1, S2, S3, S4, S5 and S6. Supplemental Table S1 and S2. for "The MacqD Deep Learning-based Model for Automatic Detection of Socially Housed Laboratory Macaques"

### Supplementary materials

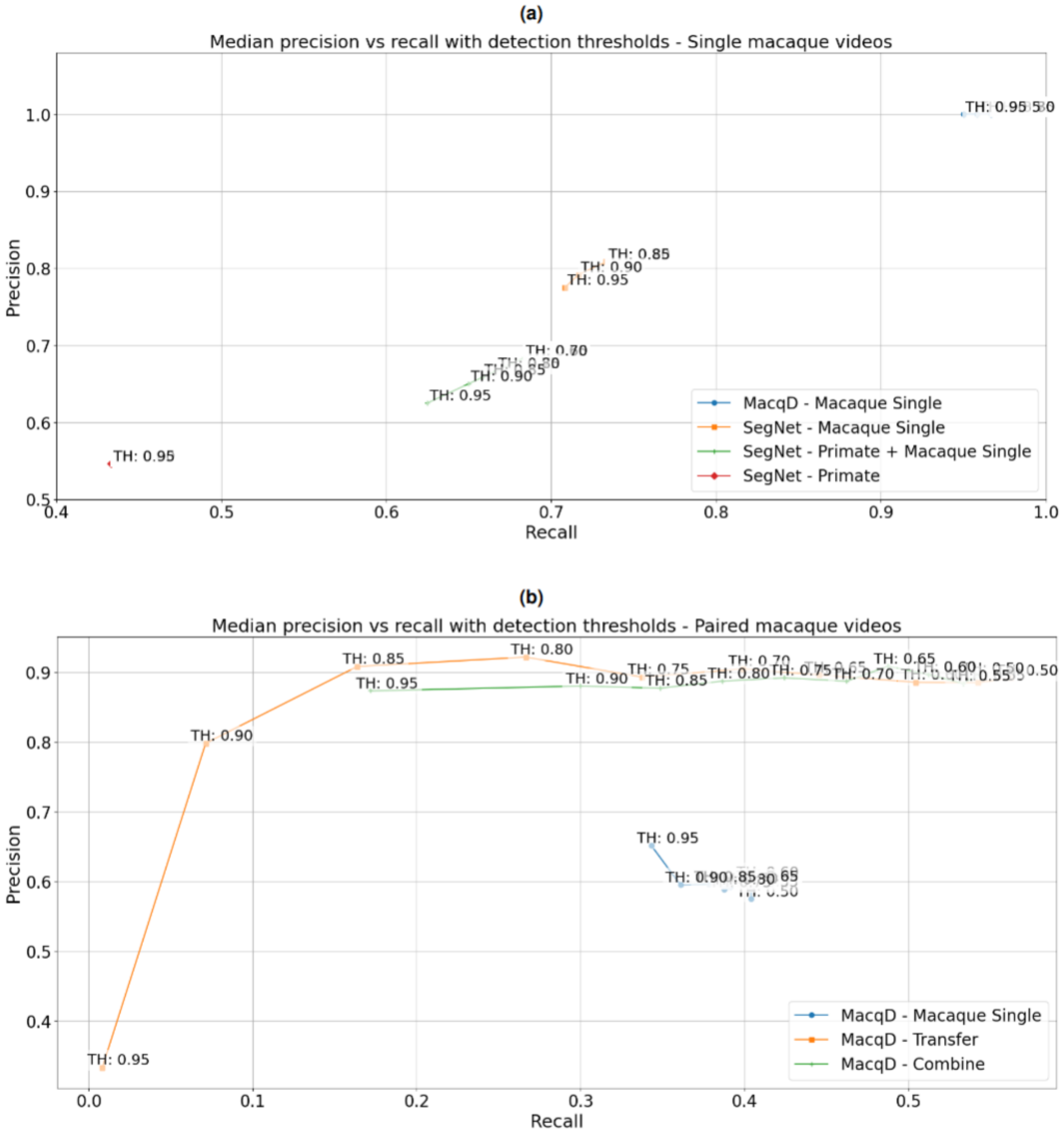


**Figure S1.** (**a**) Median precision vs. recall across various confidence score thresholds on the validation dataset with a single macaque in the focal cage. Optimal thresholds are: SegNet - Primate (0.95), SegNet - Primate + Macaque Single (0.7), SegNet - Macaque Single (0.85), and MacqD - Macaque Single (0.8). (**b**) Median precision vs. recall across various confidence score thresholds on the validation dataset with a pair of macaques in the focal cage. The optimal threshold for MacqD - Macaque Single, MacqD - Macaque Curriculum, and MacqD - Macaque Combine is 0.5.

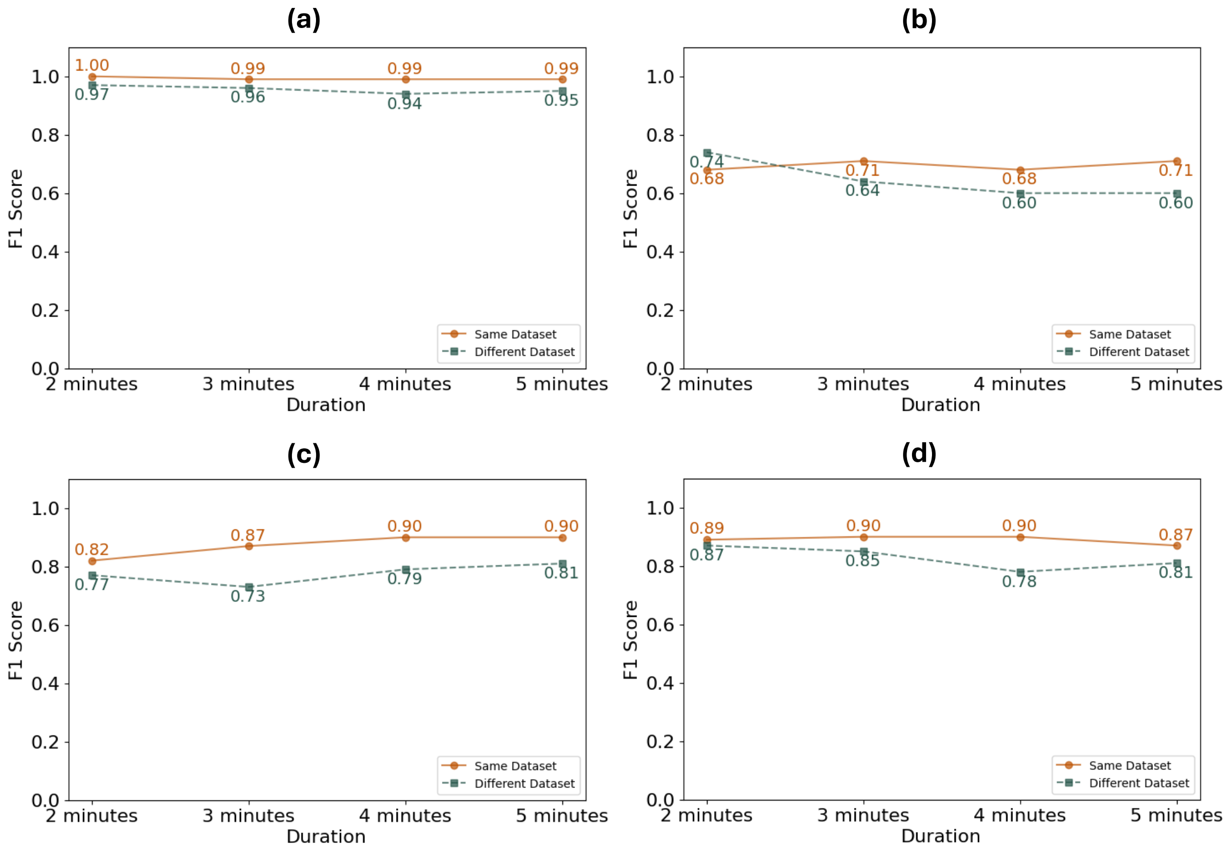


**Figure S2.** F1 scores of MacqD tested with different video durations (2–5 minutes). (**a**) MacqD - Macaque Single tested on a dataset featuring a single macaque in the focal cage. (**b**) MacqD - Macaque Single tested on a dataset featuring paired macaques in the focal cage. (**c**) MacqD - Macaque Curriculum tested on a dataset featuring paired macaques in the focal cage. (**d**) MacqD - Macaque Combine tested on a dataset featuring paired macaques in the focal cage. "Same dataset" refers to individuals that were included in the training dataset, while "Different dataset" refers to individuals that were not used during training. The F1 scores at 5 minutes correspond to the values reported in the Results section.

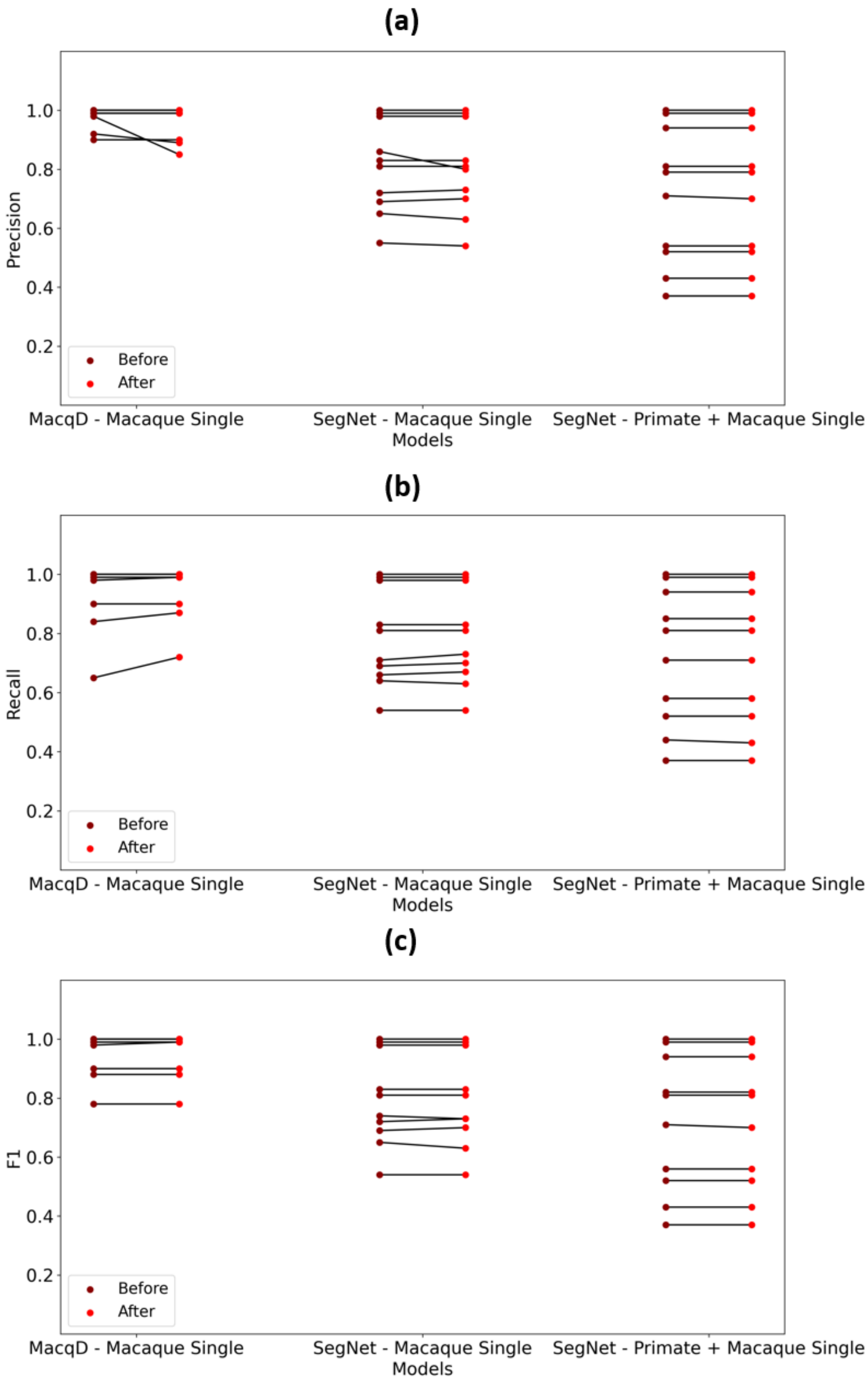


**Figure S3.** Visual representation of model predictions on frames featuring single macaques in the focal cage, which are the same individuals from the training dataset (Experiment 1, 'Same' dataset), before and after applying the tracking algorithm. (**a**) Precision; (**b**) Recall; (**c**) F1 Score.

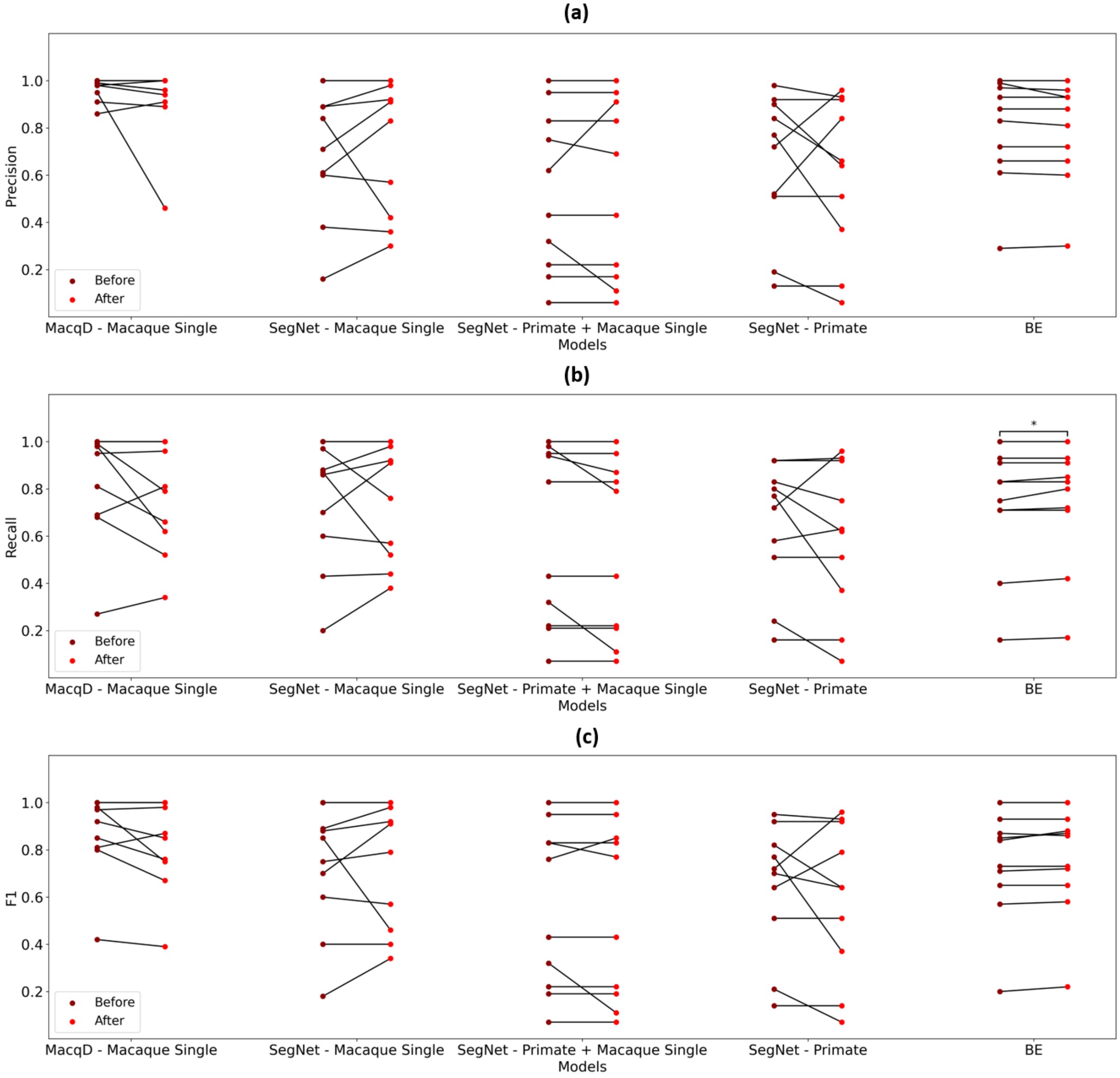


**Figure S4.** Visual representation of model predictions on frames featuring single macaques in the focal cage, which are the different individuals from the training dataset (Experiment 1, 'Different' dataset), before and after applying the tracking algorithm. (**a**) Precision; (**b**) Recall; (**c**) F1 Score. Bar indicates significant differences before and after applying the tracking algorithm (Wilcoxon test: $p < 0.05$).

**Model performance**

| Model | Dataset | Before tracking algorithm implementation: Precision | Recall | F1 | After tracking algorithm implementation: Precision | Recall | F1 |
|---|---|---|---|---|---|---|---|
| MacqD - Macaque Single | 'Same' | 0.99 | 0.99 | 0.99 | 0.99 | 0.99 | 0.99 |
| SegNet - Macaque Single | 'Same' | 0.82 | 0.76 | 0.78 | 0.81 | 0.77 | 0.77 |
| SegNet - Primate + Macaque Single | 'Same' | 0.75 | 0.76 | 0.76 | 0.75 | 0.76 | 0.76 |
| MacqD - Macaque Single | 'Different' | 0.99 | 0.97 | 0.95 | 0.96 | 0.8 | 0.86 |
| SegNet - Macaque Single | 'Different' | 0.78 | 0.87 | 0.8 | 0.87 | 0.84 | 0.85 |
| SegNet - Primate + Macaque Single | 'Different' | 0.53 | 0.63 | 0.60 | 0.56 | 0.61 | 0.6 |
| SegNet - Primate | 'Different' | 0.75 | 0.75 | 0.71 | 0.65 | 0.53 | 0.64 |
| BE | 'Different' | 0.86 | 0.79 | 0.78 | 0.85 | 0.82 | 0.80 |

Wilcoxon test

| Model | Dataset | N | DOF | Metric of comparison | $T^+$ | p-value |
|---|---|---|---|---|---|---|
| MacqD - Macaque Single | 'Same' | 10 | 1 | Precision | 0 | 0.180 |
| | | | | Recall | 0 | 0.109 |
| | | | | F1 | 0 | 0.317 |
| SegNet - Macaque Single | 'Same' | 10 | 1 | Precision | 4 | 0.336 |
| | | | | Recall | 2 | 0.257 |
| | | | | F1 | 4 | 0.705 |
| SegNet - Primate + Macaque Single | 'Same' | 10 | 1 | Precision | 0 | 0.317 |
| | | | | Recall | 0 | 0.317 |
| | | | | F1 | 0 | 0.317 |
| MacqD - Macaque Single | 'Different' | 10 | 1 | Precision | 7.5 | 0.270 |
| | | | | Recall | 6 | 0.176 |
| | | | | F1 | 4 | 0.091 |
| SegNet - Macaque Single | 'Different' | 10 | 1 | Precision | 11.5 | 0.362 |
| | | | | Recall | 16 | 0.779 |
| | | | | F1 | 8 | 0.310 |
| SegNet - Primate + Macaque Single | 'Different' | 10 | 1 | Precision | 3 | 1 |
| | | | | Recall | 0 | 0.109 |
| | | | | F1 | 2 | 0.593 |
| SegNet - Primate | 'Different' | 10 | 1 | Precision | 10 | 0.499 |
| | | | | Recall | 9 | 0.398 |
| | | | | F1 | 10 | 0.499 |
| BE | 'Different' | 10 | 1 | Precision | 2 | 0.131 |
| | | | | Recall | 0 | **0.042** |
| | | | | F1 | 2 | 0.071 |

**Table S1.** Summary of model performance and statistical comparisons on frames featuring single macaques in the focal cage (Experiment 1), before and after applying the tracking algorithm. Bold number indicates a significant difference in the metric results before and after applying the tracking algorithm.

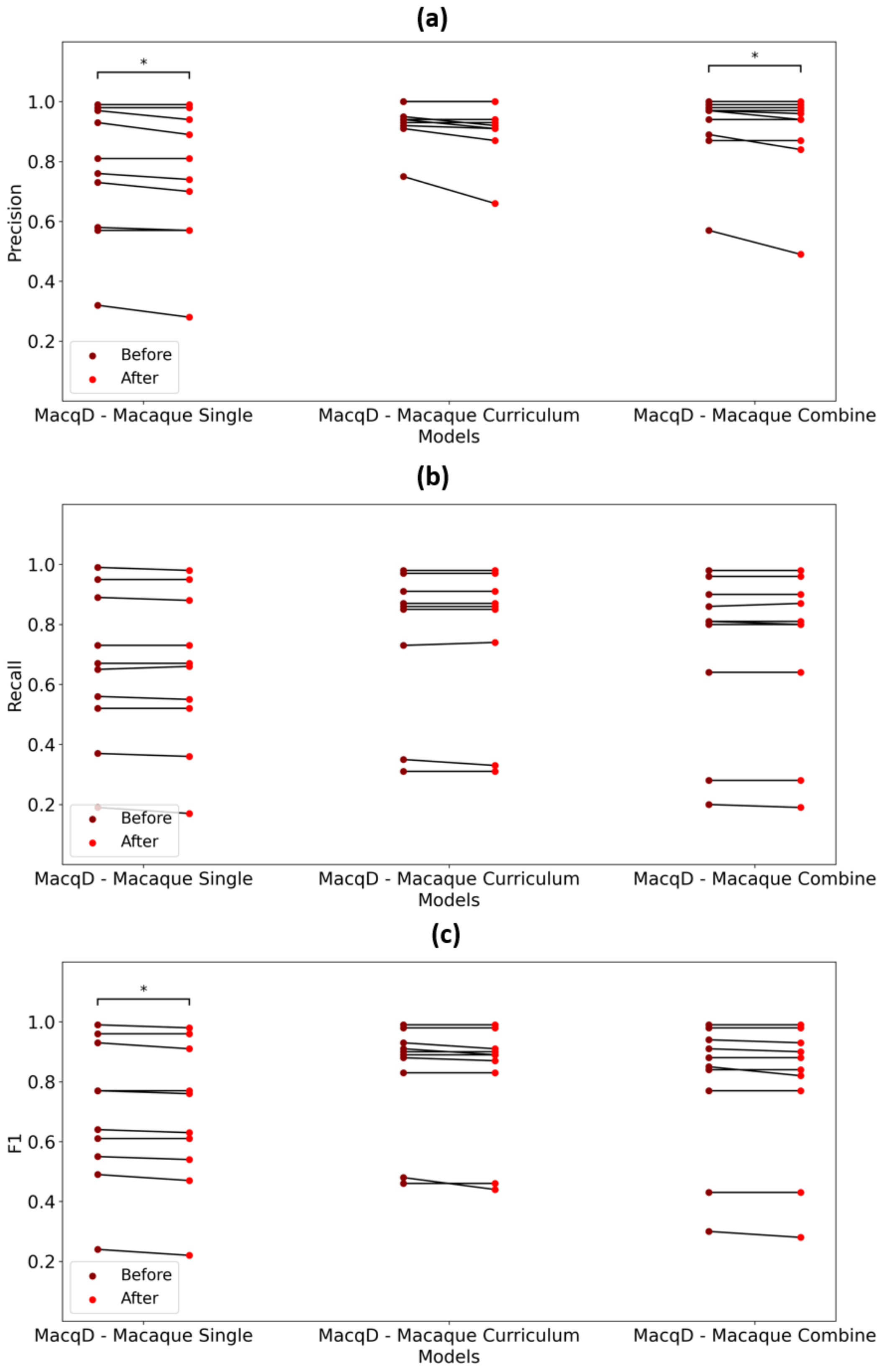


**Figure S5.** Visual representation of model predictions on frames featuring paired macaques in the focal cage, which are the same individuals from the training dataset (Experiment 2, 'Same' dataset), before and after applying the tracking algorithm. (**a**) Precision; (**b**) Recall; (**c**) F1 Score. Bars indicate significant differences before and after applying the tracking algorithm (Wilcoxon test: $p < 0.05$).

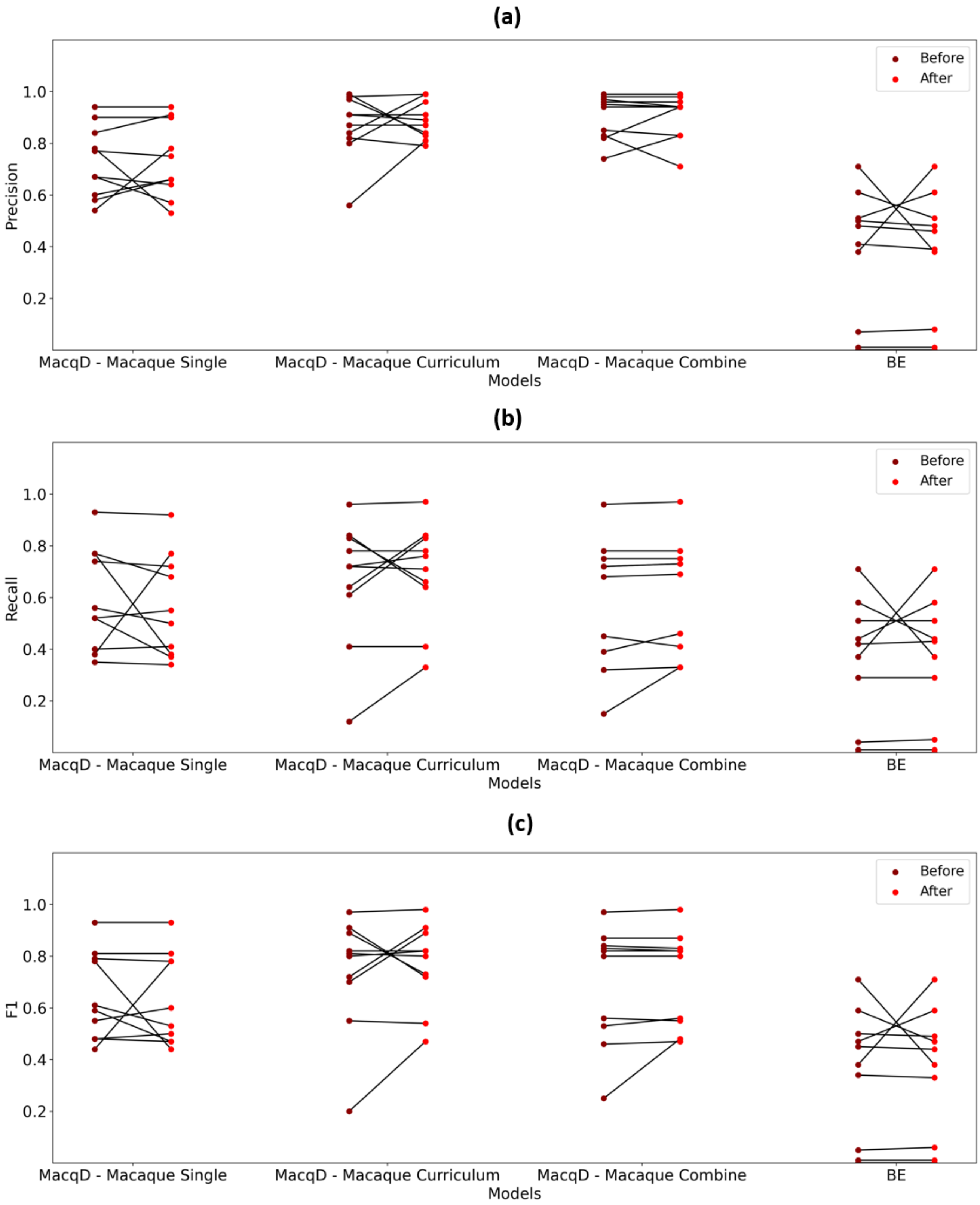


**Figure S6.** Visual representation of model predictions on frames featuring paired macaques in the focal cage, which are the different individuals from the training dataset (Experiment 2, 'Different' dataset), before and after applying the tracking algorithm. (**a**) Precision; (**b**) Recall; (**c**) F1 Score.

**Model performance**

| Model | Dataset | Before tracking algorithm implementation | | | After tracking algorithm implementation | | |
|---|---|---|---|---|---|---|---|
| | | Precision | Recall | F1 | Precision | Recall | F1 |
| MacqD - Macaque Single | 'Same' | 0.79 | 0.66 | 0.71 | 0.78 | 0.67 | 0.70 |
| MacqD - Macaque Curriculum | | 0.94 | 0.87 | 0.90 | 0.93 | 0.87 | 0.89 |
| MacqD - Macaque Combine | | 0.97 | 0.81 | 0.87 | 0.95 | 0.81 | 0.86 |
| MacqD - Macaque Single | 'Different' | 0.72 | 0.54 | 0.6 | 0.71 | 0.53 | 0.57 |
| MacqD - Macaque Curriculum | | 0.89 | 0.72 | 0.81 | 0.88 | 0.74 | 0.81 |
| MacqD - Macaque Combine | | 0.95 | 0.7 | 0.81 | 0.94 | 0.71 | 0.81 |
| BE | | 0.45 | 0.40 | 0.42 | 0.43 | 0.40 | 0.41 |

**Wilcoxon test**

| Model | Dataset | N | DOF | Metric of comparison | $T^{+}$ | p-value |
|---|---|---|---|---|---|---|
| MacqD - Macaque Single | 'Same' | 10 | 1 | Precision | 0 | **0.027** |
| | | | | Recall | 3 | 0.096 |
| | | | | F1 | 0 | **0.016** |
| MacqD - Macaque Curriculum | | | | Precision | 0 | 0.068 |
| | | | | Recall | 2 | 0.564 |
| | | | | F1 | 0 | 0.068 |
| MacqD - Macaque Combine | | | | Precision | 0 | **0.042** |
| | | | | Recall | 1 | 0.655 |
| | | | | F1 | 0 | 0.066 |
| MacqD - Macaque Single | 'Different' | | | Precision | 17 | 0.889 |
| | | | | Recall | 16 | 0.275 |
| | | | | F1 | 14.5 | 0.623 |
| MacqD - Macaque Curriculum | | | | Precision | 9.5 | 0.833 |
| | | | | Recall | 6 | 0.089 |
| | | | | F1 | 10.5 | 0.547 |
| MacqD - Macaque Combine | | | | Precision | 16 | 0.779 |
| | | | | Recall | 22 | 0.326 |
| | | | | F1 | 16 | 0.438 |
| BE | | | | Precision | 14 | 0.574 |
| | | | | Recall | 9 | 0.752 |
| | | | | F1 | 14 | 0.573 |

**Table S2.** Summary of model performance and statistical comparisons on frames featuring paired macaques in the focal cage (Experiment 2), before and after applying the tracking algorithm. Bold numbers indicate a significant difference in the metric results before and after applying the tracking algorithm.
